# Intersection of Regulatory Analysis and Signature Reversion Uncovers Therapeutic Drugs and Targets for *SETBP1*-HD

**DOI:** 10.64898/2026.05.21.726884

**Authors:** Elizabeth J. Wilk, Sasha Taluri, Tabea M. Soelter, Brittany N. Lasseigne

## Abstract

**Background:** *SETBP1* haploinsufficiency disorder (*SETBP1*-HD) is a neurodevelopmental condition characterized by developmental delay, speech apraxia, motor deficits, and autism spectrum disorder (ASD), caused by deficiency of the transcription factor (TF) SETBP1. Since current management is limited to symptomatic relief, we defined a robust consensus molecular signature for *SETBP1*-HD and prioritized drugs that converge with SETBP1 regulatory targets.

**Methods:** We performed a meta-analysis of three independent transcriptomic datasets derived from *in vitro* models of *SETBP1* deficiency. Using Robust Rank Aggregation (RRA), we overcame transcriptomic heterogeneity to establish a *SETBP1*-HD consensus signature. We integrated this signature with known and spatio-temporal-aware SETBP1 interactions to prioritize key regulatory and drug targets. Finally, we screened the LINCS L1000 drug perturbation database for compounds capable of reversing the consensus *SETBP1*-HD gene signature.

**Results:** Our regulatory analysis yielded potential, critical therapeutic windows, while our consensus analysis identified *LIN28A* (a key regulator of developmental timing) and *SPON1* ( F-spondin) as novel, robustly upregulated targets across models. *LIN28A* provides a potential mechanism for the dysregulated neurogenesis and progenitor proliferation observed in patient-derived cells, while *SPON1* links molecular dysfunction to motor and connectivity deficits seen in patients. Our *in silico* drug repurposing screens prioritized celecoxib and buspirone as top therapeutic candidates. Mechanistically, celecoxib targets the prioritized *SETBP1* regulatory target COX-2 (*PTGS2*) to mitigate neuroinflammation, while the potent neuromodulator buspirone chemically stabilizes vulnerable circuits to correct the underlying neurotransmitter imbalance driven by *SETBP1* loss.

**Limitations:** This study relies on *in silico* consensus modeling derived from cell culture and animal models. While robust statistical thresholds were applied, the therapeutic efficacy of the identified candidates on behavioral phenotypes—specifically motor deficits and speech apraxia—requires *in vivo* validation. Additionally, the specific contribution of *LIN28A* to the ASD phenotype warrants further functional characterization.

**Conclusions:** We defined a unified molecular mechanism for *SETBP1*-HD, in which dosage deficiency leads to a delayed neurogenic state driven by *LIN28A*, coupled with connectivity defects driven by *SPON1,* ultimately resulting in a destabilized excitation/inhibition balance. We identify celecoxib and buspirone as top therapeutic candidates, offering an immediate translational pathway to address the core neurodevelopmental, circuitry, and motor challenges associated with *SETBP1-*HD variants.

## Background

*SETBP1* Haploinsufficiency Disorder (*SETBP1*-HD) is a rare neurodevelopmental disorder characterized by intellectual disability, hypotonia, motor delays, and expressive speech impairment^1–5^. It is caused by *de novo* loss-of-function variants that reduce levels of the SETBP1 transcription factor (TF). *SETBP1* is a high-confidence autism spectrum disorder (ASD) risk gene^2,4^ and is highly dosage-sensitive, meaning its protein must remain within a narrow, strictly regulated range for proper function^6,7^. Distinct clinical phenotypes arise depending on how genetic variants disrupt this delicate balance. While *SETBP1*-HD results from protein deficiency, missense variants within the degron motif prevent protein degradation, causing SETBP1 accumulation and a severe, multi-systemic disorder known as Schinzel-Giedion Syndrome (SGS)^6,8–10^. Conversely, missense variants outside of the degron region cause atypical SGS or milder SETBP1-Related Disorders (*SETBP1*-RD)^4,11^ by disrupting DNA-binding, transcription, and neuronal differentiation. The functional impact of these variants seems to correlate with their distance from the degron, suggesting pathogenic mechanisms independent of protein abundance^11^. Finally, somatic variants in *SETBP1* are known drivers of and myelodysplastic and myeloproliferative neoplasms^12,13^. Ultimately, this strict intolerance to dosage and structural alterations highlights the critical role of *SETBP1* throughout development.

Currently, there are no disease-modifying therapies for *SETBP1*-HD. Patients are limited to treatments that only manage their complex multisystem symptoms^4,14,15^, highlighting the urgent need for new drug candidates. However, *SETBP1*-HD is a rare disease with widely varying presentation and severity, likely leading to underdiagnosis and difficulty in assessing disease prevalence. Limited patient numbers and disease heterogeneity exacerbate challenges in drug development^16,17^. Drug repurposing, identifying new uses for FDA-approved drugs outside of their original indications,^18^ provides an opportunity to combat some of these obstacles.

Approval for repurposed drugs surpasses the number of FDA approvals for novel-developed drugs and provides a rapid translational approach for *SETBP1*-HD^19^. Such efforts rely on a sound understanding of the disease’s pathophysiology and prioritize drugs based on their targets and off-target pathways. Previous drug studies for *SETBP1* disorders have focused on SGS^20^ or have been limited by using one specific model (e.g., *SETBP1*-KO induced pluripotent stem cells (iPSCs))^21^, likely yielding model-specific results. This variability, coupled with the nature of SETBP1 dose-sensitivity, makes it challenging to identify robust therapeutic targets applicable to the broader *SETBP1*-HD patient population.

To address these challenges, we integrated parallel computational approaches to prioritize repurposable *SETBP1*-HD therapeutics (**Additional file 1**). First, we mapped the healthy SETBP1 regulatory targets to identify potential mimetics for restoring lost interactions. Second, we derived a robust *SETBP1*-HD consensus signature by integrating transcriptome data across studies to identify consistent (i.e., model organism and dataset independent) molecular dysfunctions. Using these approaches, we identified FDA-approved candidates with pediatric safety profiles for streamlined clinical translation, including the COX-2 inhibitor celecoxib and the 5-HT1A agonist buspirone.

## Methods

### Comprehensive SETBP1 target list annotation

We created a comprehensive SETBP1 target list by querying and combining targets across 15 databases: GTRD^22^(2019 release), TRRUST^23^(v2), CollecTRI^24^(accessed May 2023), SIGNOR^25^(v3.0), the Human Transcription Factors DB^26^(accessed February 2025), TFLink^27^(accessed February 2025), IMEX^28^(accessed February 2025), TcoF^29^(v2), BioGRID^30^(2019 release), IntAct^31^(accessed February 2025), TFcheckpoint^32^(accessed February 2025), Harmonizome^33^(accessed February 2025), PathwayCommons^34^(accessed February 2025), hTFtarget^35^(accessed February 2025), and JASPAR^36^(2018 release). Further, we manually curated literature-supported target and interaction details and inferred interaction effects for unknown target interactions based on top co-expression with *SETBP1* across over 1 million gene expression signatures, derived from LINCS^37^(2020 release) and GEO data^38^ **(Additional file 2)**.

### SETBP1 target spatio-temporal permutation testing and prioritization

We then validated SETBP1 targets exhibiting temporally appropriate developmental reorganization (i.e., targets expressed at specific developmental times) with permutation testing using BrainSpan (developmental transcriptome, RNA-seq summarized to genes; accessed March 2026)^39,40^, dGTEX, and GTEx (RNA-seq transcripts per million [TPM], v1 and v11, respectively)^41,42^. We log2-transformed, filtered for non-zero-variance genes, and scaled expression counts. For BrainSpan, we categorized samples by developmental windows: Early Prenatal (≤ 12 post-conceptional weeks [pcw]), Mid Prenatal (13-24 pcw), Late Prenatal (25-40 pcw), Infancy (≤ 1 year), Early Childhood (2-5 years), Late Childhood (6-17 years), and Adulthood (≥ 18 years). Our developmental window selections were guided by synaptic milestones (i.e., rapid synaptogenesis until age 2, followed by synaptic pruning) ^43,44^ and trimesters, while also optimizing for sample number (e.g., due to limited second trimester data, we started “Mid Prenatal” range at 13 pcw). For dGTEx (age brackets ranging from 0-18 years old), we combined ages as “Pediatric” due to limited samples across ages, and compared to adult GTEx age bins (age brackets in 10-year increments ranging 20-79). We assessed enrichment of SETBP1 regulatory targets within each developmental window by performing a permutation analysis. For each window, we calculated the observed test statistic as the mean scaled expression of the targets. We calculated a null distribution using the mean expression of randomly sampled background genes of an equivalent size (*n* = targets) across 10,000 permutations. We calculated two-sided p-values and Z-scores (enrichment scores) by comparing the observed target mean to the null distribution. We corrected for multiple hypotheses using Benjamini-Hochberg False Discovery Rate (FDR)^45^ p-value adjustment. To check for sex-bias, we split each tissue and age bracket by sex and ran Wilcoxon rank sum tests with FDR correction for GTEx data.

We prioritized SETBP1 regulatory targets for potential drug targeting by filtering for PHAROS^46^(accessed February 2025) target development level of Tchem or Tclin (targets with strong small molecule binding affinity or approved drugs, respectively) and by top co-expression or core pathway involvement. We defined top coexpressed targets as those with top quartile *SETBP1* co-expression across our spatio-temporal conditions in BrainSpan and GTEx data. For assessing core pathway involvement, we mapped targets to Pan-GO^47^ categories consistent with three previously documented overarching SETBP1 mechanisms: 1) transcriptional and epigenetic regulation^7,48,49^, 2) cell proliferation and survival^6,7^, and 3) neurodevelopment and circuitry^7,50^.

### RNA-seq data set curation, alignment, and quantification

We obtained raw RNA-seq reads using the SRA Toolkit (v3.2.1)^51^ from three independent SETBP1 deficiency studies (**Additional file 3**): Cardo et al. *SETBP1*-/- human embryonic stem cells (hESCs) neural progenitor cells (NPCs) and neurons (harvested at days 15, 21, and 34)^21^, Shaw et al. induced pluripotent stem cells (iPSCs) and NPCS engineered with *SETBP1* LoF truncation variants (p.Glu545Ter and p.Tyr1066Ter) and a VUS (p.Thr1387Met)^50^, and Tanaka et al. hematopoietic stem/progenitor cells from Vav1-iCre;*Setbp1^fl^*^/fl^ conditional knockout mice^52^. For quality control, alignment, and quantification, we used the Nextflow nf-core/rnaseq pipeline (v3.18.0)^53,54^. We aligned human data to the GRCh38 genome (GENCODE v47 annotation), and mouse data to the GRCm39 genome (GENCODE v31 annotation)^55,56^.

### Exploratory data analysis

We cleaned and prepared gene count data by filtering out genes with zero variance, then explored sample-sample correlations with respect to metadata (tidyverse v2.0.0)^57^ and performed principal components analysis (PCA) using the top 10,000 most variable genes (base R 4.3.1). We built random forest (ranger v0.17.0)^58^ and SVM (e1071 1.7-16)^59^ models with XAI (DALEX v2.5.2)^60^ to predict metadata features driving principal components and plotted results (**Additional file 4**).

### Differential expression analysis

To find differentially expressed genes (DEGs), we first filtered low-count genes (filterByExpr()), normalized (calcNormFactors()), and mean-variance transformed (voom()) using limma-voom (limma v3.56.2, edgeR v3.42.4)^61,62^. We applied linear modeling and empirical bayes (limma v3.56.2) lmFit(), contrasts.fit(), eBayes(), and computed DEGs for each contrast using topTable() with BH multiple hypothesis correction. We retained DEGs |logFC| > 0.5 and adj p-val < 0.1, and calculated standard error (absolute value of the logFC/t-statistic).

### Consensus gene expression signature

To derive a robust consensus signature, we tested multiple analytical methods: frequency-based, meta-analysis, aggregate ranks, and composite score-based. We first identified DEGs common to at least one contrast (i.e., *SETBP1* KO vs NPC at differentiation day 21) in both human *SETBP1*-deficiency datasets (Cardo et al. and Shaw et al.). High-frequency genes (freq = 5) were identified by counting occurrences across heterogeneous contrasts, including multiple developmental time points and genotypes.

We performed an effect size meta-analysis, using the metafor R package (v4.8)^63^, fitting a linear model via restricted maximum likelihood (REML) based on each gene’s logFC and standard error. Additionally, we found aggregate ranks via RobustRankAggregate (v1.2.1)^64^ to identify genes consistently ranked across contrasts. To compute composite scores, we combined results from the three signature methods:

*Composite score* = -log10(*meta-analysis p-val*) + -log10(*RRA p-val*) + log2(*n*)

To ensure compatibility with the LINCS drug reference databases, we mapped Ensembl gene IDs to Entrez IDs. We defined candidate signatures by selecting the top 300 genes (based on prior work^65^) for each signature method (frequency, meta-analysis p-value, rank aggregate p- value, and composite score). We applied over-representation analysis (ORA) to each signature (gprofiler2 v0.2.3)^66^ using GO, KEGG, and WikiPathway sources with terms with < 400 genes to exclude broad parent terms.

We validated signatures against orthogonal DEGs from Tanaka et al. and Wong et al., as well as high-confidence *SETBP1*-associated pathways, including forebrain development (GO:0030900), PI3K/AKT signaling (REAC:R-HSA-5653656), and Wnt signaling (KEGG:04310)^6,11,21,50,67^. To do this, we evaluated signature performance using precision-recall and F1-scoring, treating orthogonal DEGs and pathway-specific genes as “true positives” against the background of all combined Cardo and Shaw DEGs. We obtained Wong et al. pre-calculated DEGs from their Source data (Source Data 6 and Source Data 10) and accompanying metadata (Supplementary Data 5 and Supplementary Data 7) for patient-derived fibroblasts and induced neurons, respectively.

### Signature reversion drug analysis

We mapped our RRA consensus signature to Entrez Gene IDs in the LINCS L1000^68^ reference database and filtered the results for the top 300 genes (by p-value for meta-analysis and RRA) to obtain ∼100 up- and down-mappable genes (based on prior work^65^). We queried the LINCS compound-treated cell database (2020 release) using the GESS algorithm with the signatureSearch^69^ R package and calculated reversion metrics (i.e., weighted connectivity scores [WTCS]) to identify signatures that would “reverse” our consensus *SETBP1*-HD signature. To focus on the most relevant cell line model for our study, we filtered the resulting LINCS signatures for brain/CNS-derived cells (NEU, SHSY5Y, and NPC) and for highly similar cells often used in neuroscience studies (HEK293 and HEK293T)^70–72^. We annotated drugs’ target genes, mechanisms of action (MOAs), and indications with signatureSearchData data (lincs_pert_info2)^69^, and annotated safety data using DailyMed API (accessed 260219, data version download available on Zenodo (10.5281/zenodo.20073644)^73^).

### Visualizations

Additional file 1, graphical abstract was made in Biorender. All other visualizations were made in R using ComplexHeatmap (v2.16.0)^74^, UpSetR (v1.4.0)^75^, and ggplot2 (v4.0.0)^76^.

### Code and Reproducibility

All code was built using the CAPTURE framework (cap version 1.0)^77^. To ensure rigor and reproducibility, all code was reviewed and independently tested by two independent lab members (i.e., not the code author) and by using docker images (see “Availability of Data and Materials” for more)^78^. All code to reproduce analyses are available at: https://github.com/lasseignelab/setbp1_hd/tree/main and (10.5281/zenodo.20073553)^79^ .

## Results

### Spatio-Temporal Expression and Functional Prioritization Identifies Key Neurodevelopmental Targets of SETBP1

As *SETBP1*-HD is fundamentally caused by a lack of the SETBP1 TF, we hypothesized that the restoration of context-specific SETBP1 interactions may prevent and/or treat the disorder (**Additional file 1**). We began with curating a comprehensive table of 361 known and predicted regulatory targets of SETBP1 across databases and manual literature review (see Methods). Of note, our manual review uncovered an interaction misnomer–*PTGS2* and *PTGER2* were indicated as repressed by SETBP1–due to a misciting of a paper on Staphylococcus aureus enterotoxin B (SEB; also a synonymous name for *SETBP1*)^80^. Some evidence corroborates *PTGS2* as a target, finding upregulated expression in SGS^81^, suggesting SETBP1 activation/stabilization rather than repression. Therefore we annotated these interactions accordingly.

To identify critical spatio-temporal contexts for SETBP1 regulation, we compared our SETBP1 target genes’ expression to a null set of genes in BrainSpan and GTEx data. Across prenatal and postnatal brain development, we observed significantly different enrichment of targets compared to null gene sets within BrainSpan RNA-seq data (**Figure 1A**). SETBP1 targets are the most enriched during early prenatal brain development (8-12 pcw, z-score = 9.51), during rapid NPC, neuronal, and macroglial expansion^43,44^. Conversely, SETBP1 targets are the most depleted during early childhood (2-5 years, z-score = -9.24), a phase marked by synaptic pruning and myelination^44^, highlighting potential critical windows of SETBP1 regulation.

**Figure 1:**
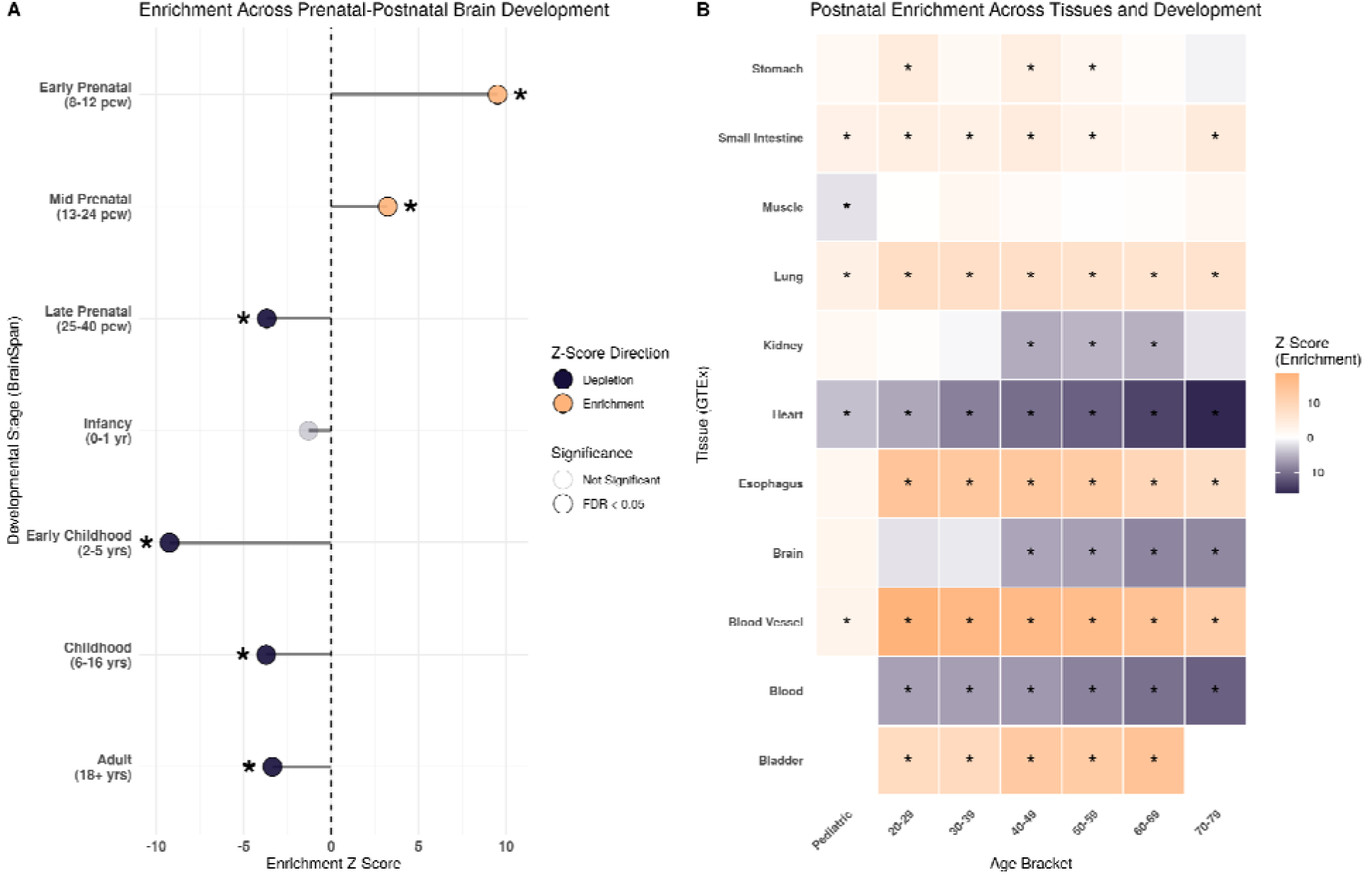
SETBP1 Targets Spatio-Temporal Permutation Analysis. Permutation analysis results for the SETBP1 target gene set compared to null gene sets across development in A) prenatal and postnatal human brain and B) postnatal *SETBP1* disorder-associated tissues.

We further investigated context specificity through postnatal enrichment across 11 tissues with phenotypes across *SETBP1*-associated disorders^82^. In adult tissues (ages 20-79), SETBP1 targets were robustly depleted in blood and heart (z-score < 0, p-adj < 0.05), while significantly enriched in the blood vessel, esophagus, lung, and small intestine (z-score > 0, p-adj < 0.05) (**Figure 1B**). Aggregated pediatric timepoints (ages 0-18 years) largely reflected these trends by tissue (i.e., same direction regarding enrichment/depletion), with the exception of muscle, kidney, and brain (**Figure 1B**). Although changes were mostly insignificant across ages, pediatric kidney and brain had slight enrichment (z-scores 1.65 and 2.24, p-adj 0.14 and 0.05, respectively), contrary to the depletion observed in the adult kidney and brain tissues. However, due to currently limited sample numbers in dGTEx, we were underpowered to detect significant enrichment in pediatric brain and kidney (n=4 and n=3, respectively). Interestingly, we found significant pediatric-specific depletion of SETBP1 targets in muscle (z-score = -1.97, p-adj = 0.049), compared to slight, insignificant enrichment across the adult age brackets (z-score range 0.08 - 1.97, p-adj range 0.07 - 0.92). As there are sex differences in ASD prevalence and presentation, we also compared SETBP1 target expression by sex across conditions in GTEx, but did not observe any significant differences (**Additional file 5**).

For downstream drug prioritization, we filtered for regulatory targets with known chemical and/or drug binding (PHAROS Tchem or Tclin, respectively) and with top *SETBP1* spatio-temporal co-expression or core *SETBP1* pathway involvement (i.e., 1) transcriptional and epigenetic regulation^7,48,49^, 2) cell proliferation and survival^6,7^, and 3) neurodevelopment and circuitry^7,50^). Our prioritization schema resulted in 34 actionable targets with high SETBP1 association (by co-expression and/or core pathway involvement) (**Table 1**). While these targets offer potentially relevant spatio-temporal interactions in non-diseased conditions, they may not recapitulate perturbations within the disease state.

**Table 1:**
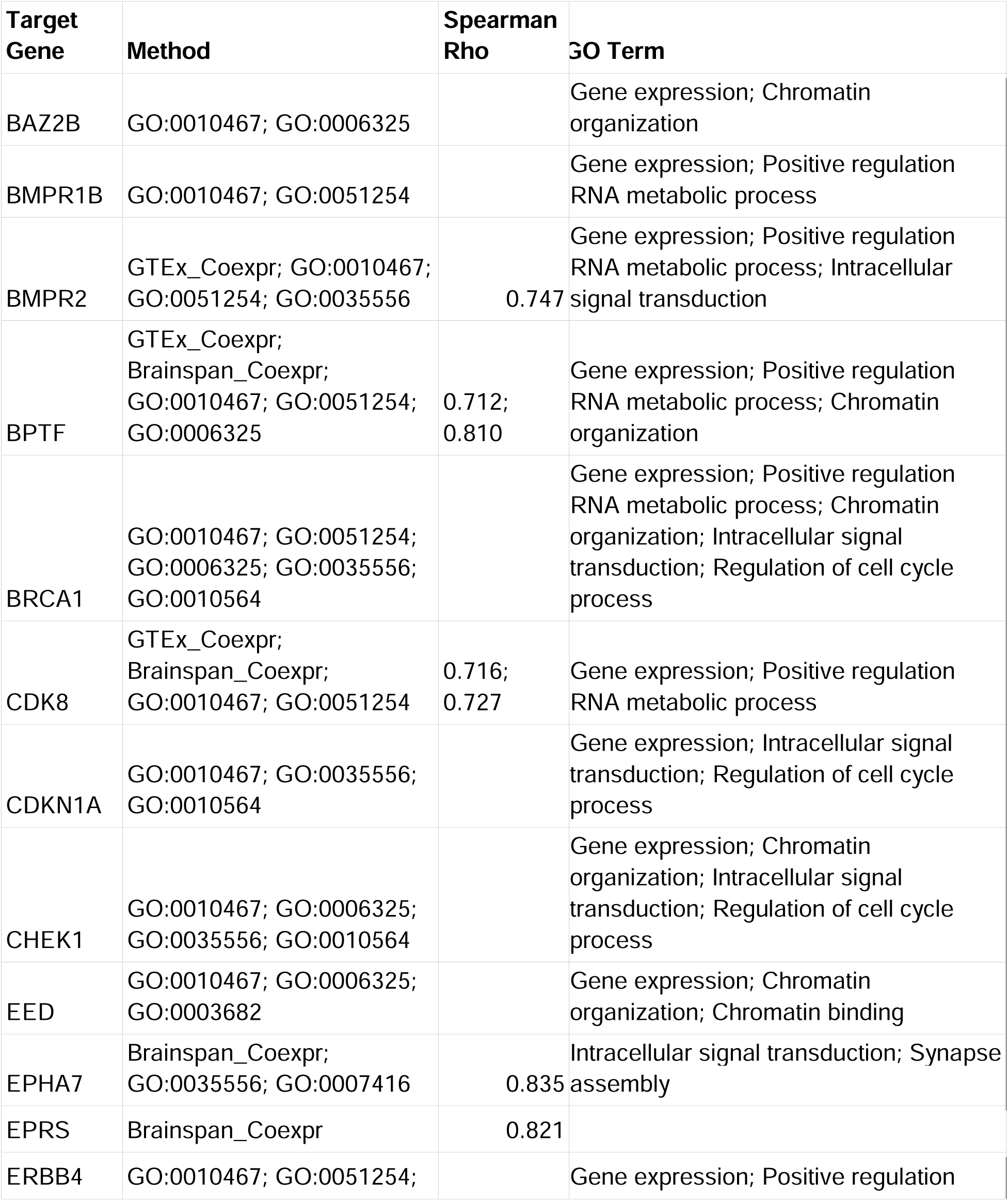
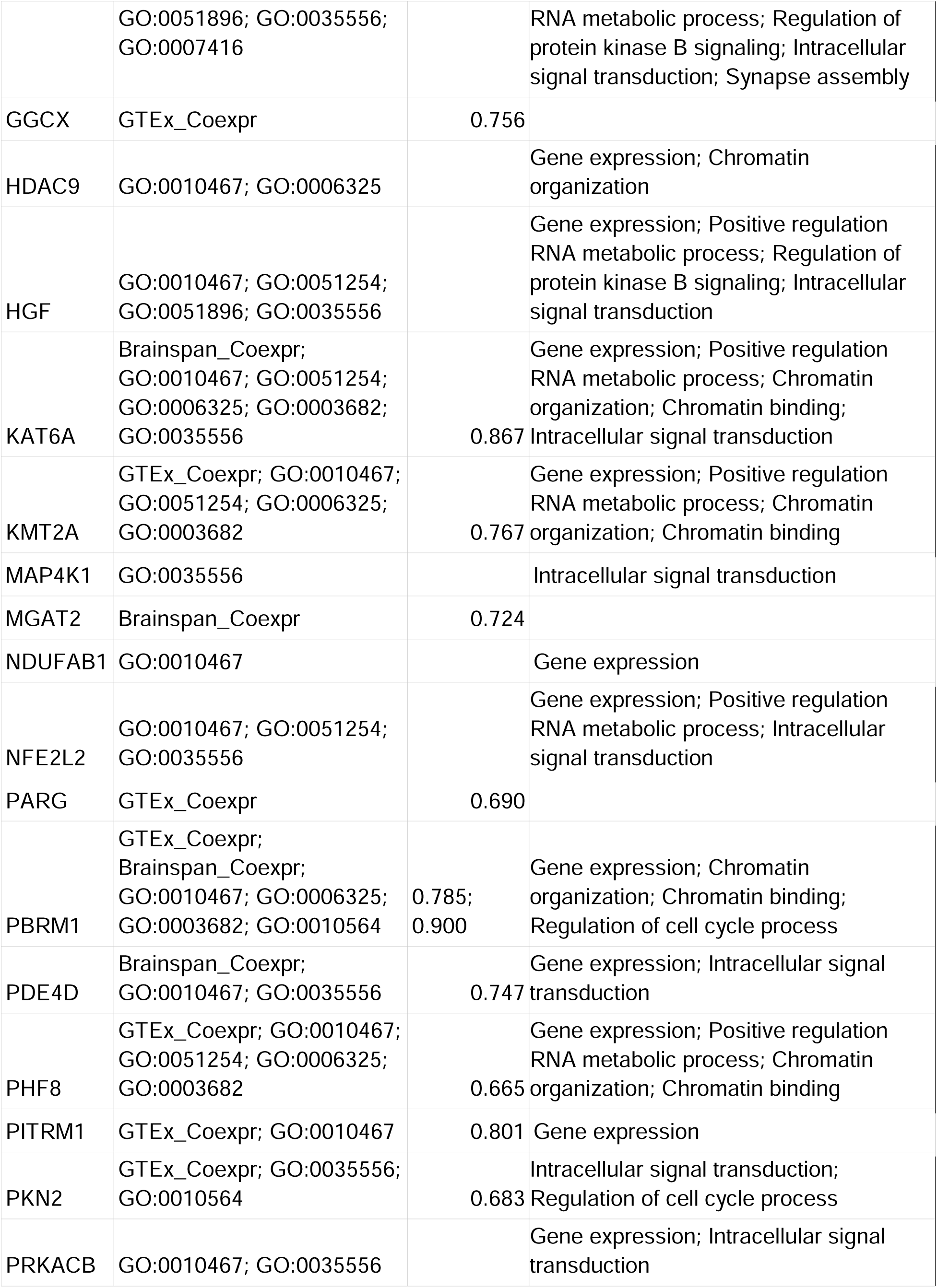
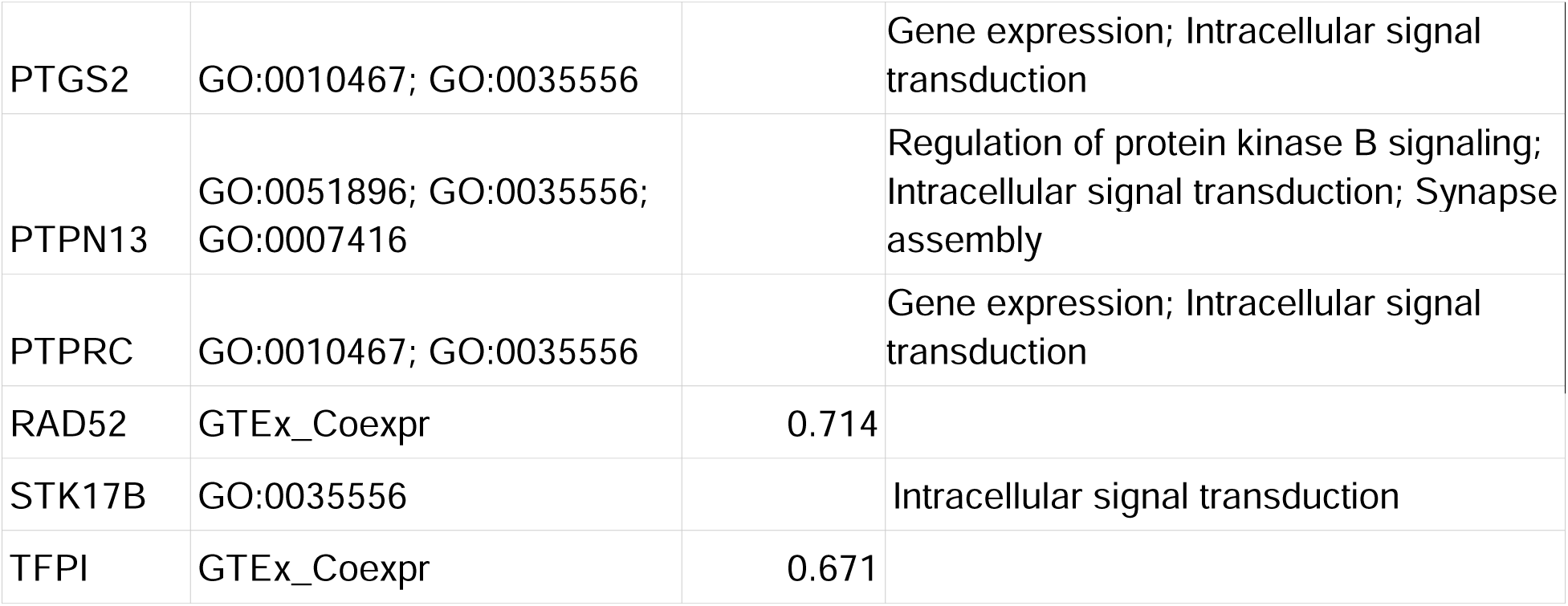
Prioritized SETBP1 regulatory targets. Prioritized SETBP1 regulatory targets. Targets were prioritized for potential drug repurposing based on known druggability and strong biological association. Association was determined via top-quartile spatio-temporal co-expression with *SETBP1* (GTEx and BrainSpan databases) and/or involvement in core *SETBP1*-associated Gene Ontology (GO) pathways, such as transcriptional regulation, cell proliferation, and neurodevelopment. The table details the inclusion method, Spearman correlation (Rho) for co-expressed targets, and the specific GO IDs and terms for each gene.

### Meta-Analytic Robust Rank Aggregation Overcomes Dataset Heterogeneity to Define a Robust *SETBP1*-HD Consensus Signature

To confirm that SETBP1 interactions are perturbed across diverse biological contexts (genotypes and differentiation stages), we curated two public *SETBP1*-HD data sets (Cardo et al. and Shaw et al.). Cardo et al. used CRISPR *SETBP1*-KO hESCs to study deficiency across neurogenesis time points (differentiation days 15, 21, 34)^21^. Shaw et al. focused on variant-specific perturbations using two pathogenic patient variants (p.Glu545Ter “PATH2” and p.Tyr1066Ter “PATH3”) and one variant of unknown significance (VUS) (p.Thr1387Met)^50^.

We compared these human data sets to an orthogonal mouse data set (Tanaka)^52^, and found that organism (human vs. mouse) accounted for the greatest variation (PC1 27.77%), followed by cell type (PC2 22.65%) (**Additional file 4**). Notably, we did not observe any PCs (total variation > 3%) driven by control vs. *SETBP1*-deficient samples, likely reflecting the heterogeneity observed across *SETBP1*-HD patients and models.

To overcome this heterogeneity across models–in hopes of overcoming patient heterogeneity, as well–we prioritized liberal thresholds to detect smaller effect sizes (|logFC| > 0.5, adjusted p-value < 0.1). Consistent with Shaw et al., we observed large differences between the two pathogenic variants: PATH2 (p.Glu545Ter) showed significant differential expression (4,279 DEGs), whereas PATH3 (p.Tyr1066Ter) showed almost no differential expression compared to wildtype^50^ (6 DEGs) (**Additional file 7**). Consequently, we designed our differential expression contrasts by individual genotype and timepoint (VUSvsWT, PATH2vsWT, and PATH3vsWT) rather than pooling all *SETBP1* genotypes.

To identify a consensus, we filtered for genes differentially expressed in at least one contrast across datasets. The Cardo dataset showed high internal overlap across contrasts (**Additional file 8**), likely due to the strong phenotype of the complete KO model compared to the patient-variant models in Shaw. While we did not find any overlapping DEGs between Shaw PATH3 and Cardo, we identified 626 DEGs shared between Shaw PATH2 and Cardo (all timepoints), and 306 DEGs shared between Shaw PATH2, Shaw VUS, and Cardo (all timepoints; **Additional file 7**).

We tested multiple analytical methods to overcome heterogeneity and derive a robust, representative, and biologically relevant consensus signature. We compared frequency-based, effect size meta-analysis, robust rank aggregation (RRA), and composite score methods, followed by validation with orthologous datasets and functionally-relevant *SETBP1*-HD genes (i.e., genes involved in Wnt and neurodevelopment pathways). We filtered the results from each method for the top 300 genes (by p-value for meta-analysis and RRA) to obtain ∼100 up- and down-mappable genes for downstream signature-reversion analyses (based on prior work^65^).

We performed ORA for each of these signatures using the GO, KEGG, and WikiPathway gene sets. This analysis showed the greatest number of enriched terms for frequency-based and RRA-based signatures (**Figure 2A**), suggesting that these methods captured more biologically relevant signals. Additionally, terms enriched for the RRA signature aligned particularly well with established disease mechanisms (**Figure 2A**), including ‘neuron projection guidance’^6^, ‘regulation of neuron differentiation’^21,50^, ‘neural precursor cell proliferation’^21^, and ‘telencephalon cell migration/development’^21^.

**Figure 2:**
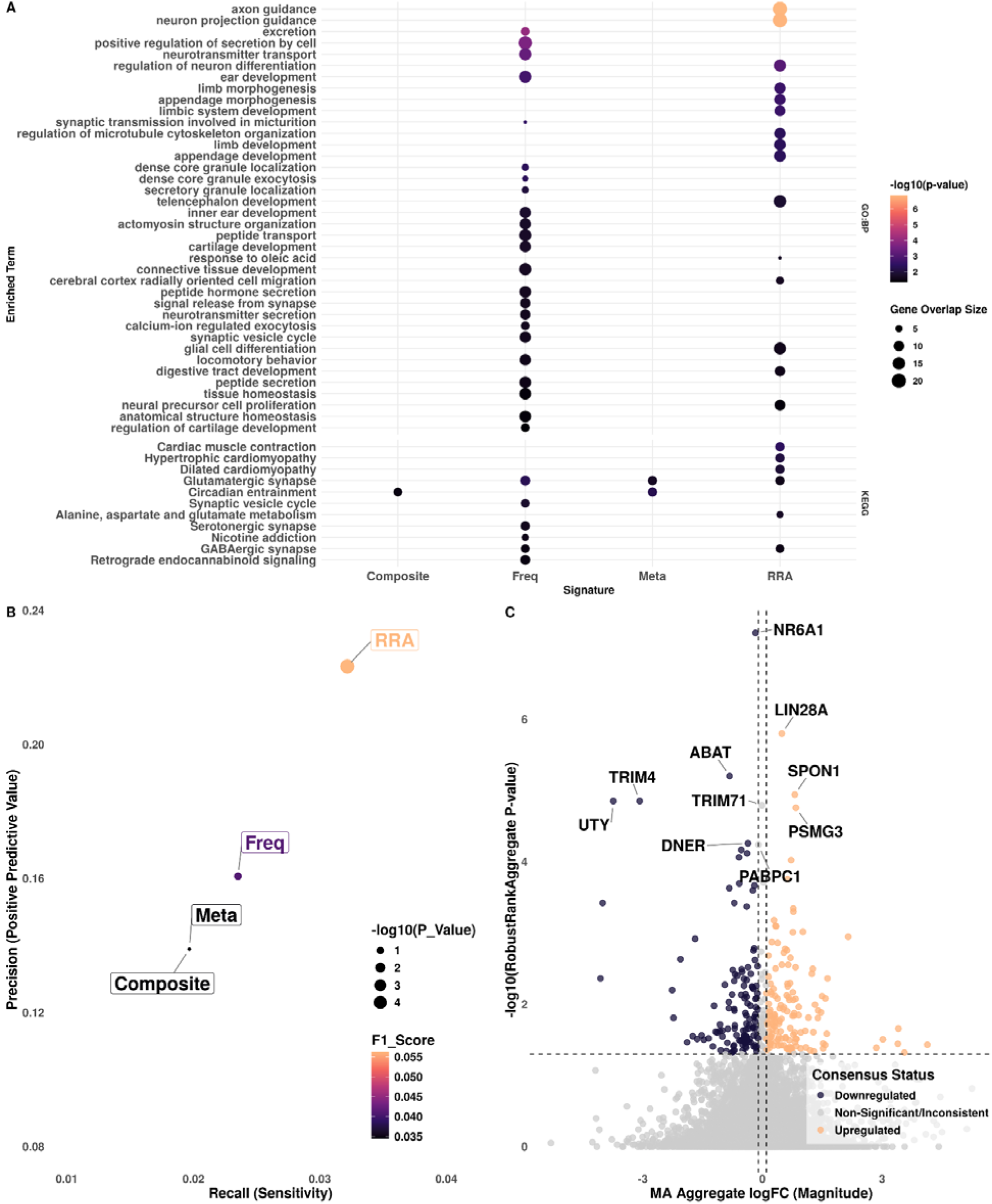
RRA-Based Consensus Signature Captures Known *SETBP1* Transcriptomic and Biological Mechanisms. Consensus method comparisons of composite-, frequency-, meta-analysis, and RRA-based signatures. A) Bubbleplot of ORA terms (geneset terms filtered for < 400 genes) enriched for each signature method, with size indicating the gene overlap (number of genes shared between signature and term geneset) and -log10 adjusted p-value from ORA. B) Dotplot of signature performance by method (RRA, Freq, Meta, and Composite) by Recall (x) and Precision (y), with size indicating -log10(p-value) and colored by F1 score. C) Volcano plot of RRA -log10 adjusted p-values (y axis) by meta-analysis logFCs (x axis) with RRA consensus signature colored red (upregulated RRA signature) and blue (downregulated RRA signature).

For validation, we used genes from highly supported *SETBP1*-HD-associated pathways and processed DEGs from Tanaka et al. and Wong et al. *SETBP1* truncation-variant samples^11,52^. We compared each of our signatures to this validation gene set and found the highest overlap with RRA signature genes (67 shared genes, compared to 49, 41, and 41 for frequency-based, meta-analysis, and composite score signatures, respectively) (**Figure 2B**). Accordingly, we proceeded with the RRA signature and used our meta-analysis logFCs (|logFC| > 0.1) to assign directionality, thus deriving our consensus signature (**Figure 2C**).

Our top ten consensus signature genes included three upregulated genes (*LIN28A, SPON1*, and *PSMG3*), 5 downregulated genes (*NR6A1, ABAT, TRIM4, UTY*, and *DNER*), and 2 with insignificant meta-analysis logFCs (*TRIM71and PABPC1*). We found a total of 38 genes highly ranked for adjusted p-values across data sets, but with inconsistent directions or with small effect sizes (e.g., *TRIM17* and *PABPC1*; **Additional file 9**). Notably, two Wnt-associated genes (*FZD6* and *ZNRF3*) displayed conflicting directionality: upregulation in Cardo (differentiation days 21 and 34)^21^, but downregulation in Shaw (PATH2 and VUS models)^50^. These discrepancies likely reflect the broader conflict in the literature, where Cardo et al. reported Wnt/β-catenin hyperactivation in *SETBP1*-deficient models^21^, while Antonyan et al. reported decreased β-catenin levels in *SETBP1*-HD contexts^6^. While this heterogeneity may recapitulate the variability observed among *SETBP1*-HD patients, we prioritized maximizing consistency to uncover therapeutics with the broadest potential applicability.

### Signature Reversion and Target Intersection Converge on Repurposable Therapeutic Candidates

To identify drug repurposing candidates with the potential to mitigate *SETBP1*-HD molecular disease mechanisms, we reversed our consensus signature and queried the LINCS drug perturbation database in order to identify drugs with signatures predicted to reverse the consensus signature^69^. The LINCS L1000 reference database is a high-throughput transcriptomic resource that measures gene expression changes in response to different perturbations, allowing researchers to connect drugs, genes, and diseases based on shared cellular signatures^68^. From this analysis, we identified 1,091 compounds, including 136 approved drugs (FDA or international equivalents) with the potential to neutralize *SETBP1*-HD disease-associated gene expression (**Additional file 6**). These drugs represented various MOAs, with the top categories being dopamine receptor antagonist, glucocorticoid receptor agonist, and adrenergic receptor antagonist (accounting for 8, 7, and 6 drugs’ MOAs, respectively) (**Figure 3A**). The prominence of neuromodulators directly aligns with our consensus signature’s enrichment for GABAergic and glutamatergic synapse pathways (**Figure 2A**), suggesting these signature reversion candidates may compensate for underlying neurotransmitter imbalances.

**Figure 3:**
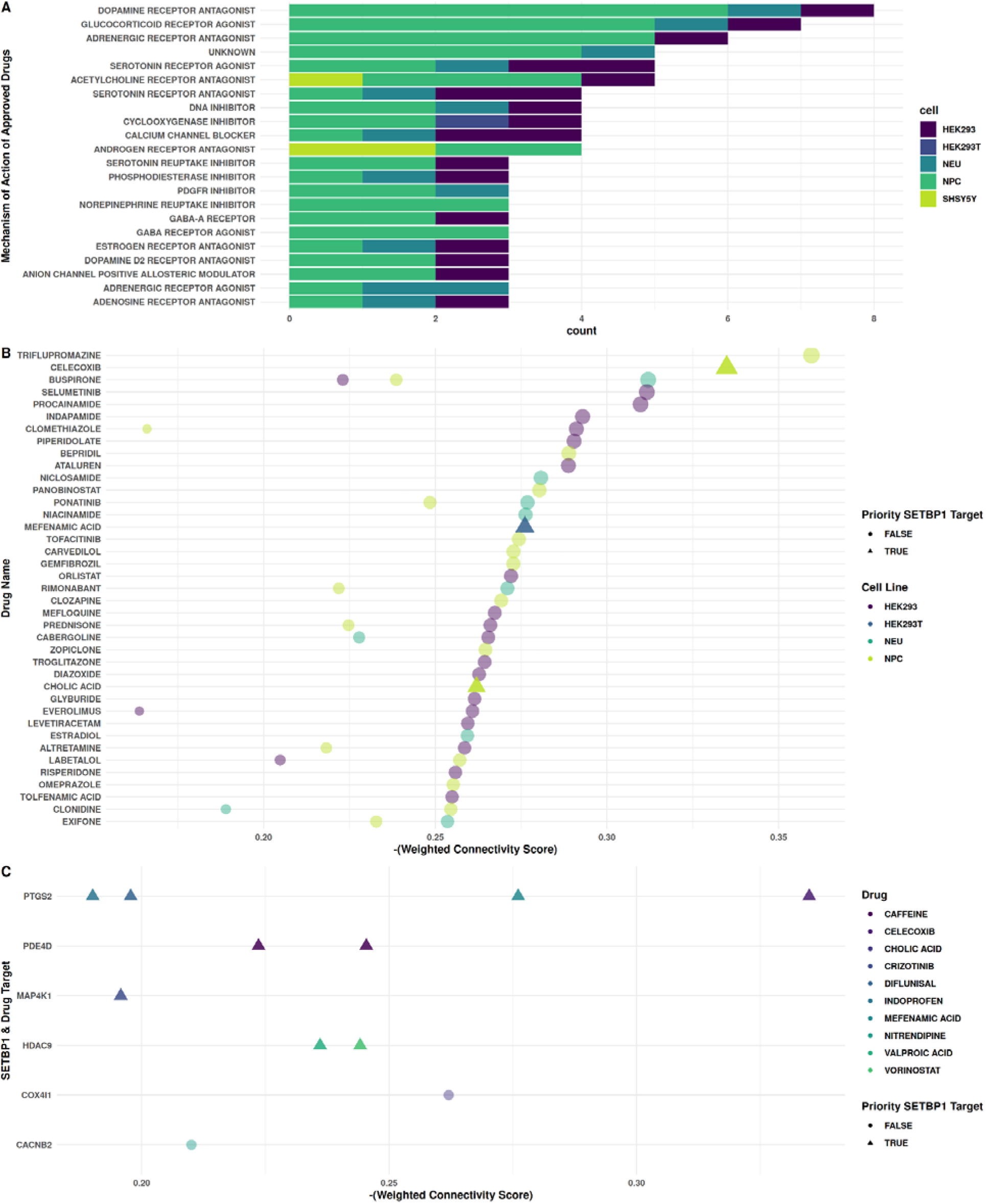
Signature reversion and regulatory network intersection prioritize *SETBP1*-HD repurposable drug candidates. Drug candidates by signature reversion. A) Frequency of Mechanisms of Action (MOAs) among the 136 identified approved drugs. Only MOAs represented by > 2 drugs are shown. B) Top 50 approved signature reversion drugs by Weighted Connectivity Score (WTCS) in brain-relevant cell lines (NEU, SHSY5Y, NPC, HEK293). Shape and transparency indicate drugs that are known to target an established SETBP1 regulatory target.C) Convergence of top-down (signature reversion) and bottom-up (regulatory network) analyses. The plot highlights 21 drug candidates that specifically act upon SETBP1 regulatory targets (y-axis)

Given that *SETBP1* is critical during prenatal development, we assessed pregnancy and pediatric safety profiles of our top 10 drugs by -WTCS to evaluate their potential for early intervention. While pregnancy safety data in humans remains limited, animal studies for several top candidates–including buspirone, indapamide, and pramipexole–have shown no evidence of fetal harm or fertility impairment at clinically relevant doses (**Additional file 6**). Additionally, celecoxib and buspirone have established profiles indicating safety in pediatric patients (ages 2 and older and ages 6 and older, respectively). The latter effectively illustrates a therapeutic potential of targeting the neurotransmitter imbalances identified in our signature **(Figure 2).** Specifically, buspirone is a partial 5-HT1A agonist and D2 antagonist that has been used to treat phenotypes that overlap with *SETBP1*-HD, including anxiety, ADHD, and cognitive impairment^83,84^.

We then evaluated whether any of our drug candidates targeted our SETBP1 regulatory targets and prioritized targets (**Additional file 2**; **Table 1**). The intersection of these “top-down” (signature reversion) and “bottom-up” (target prioritization) approaches yielded 10 unique drug candidates, linked to 6 SETBP1 targets (**Figure 3C**; **Table 2**). Of these, 4 targeted our prioritized SETBP1 targets (*PTGS2, PDE4D, MAP4K1,* and *HDAC9*), highlighting their clinical relevance across functional and spatio-temporal contexts.

**Table 2:** Convergent drug repurposing candidates. Top candidates identified by intersecting transcriptomic signature reversion results with SETBP1 regulatory targets. The table details the brain-relevant cell line, the Weighted Connectivity Score (WTCS) and p-value indicating the strength of the signature reversion, the specific SETBP1 regulatory target, and whether that target was prioritized based on spatio-temporal expression and pathway involvement.

| Name | Cell line | WTCS | WTCS Pval | SETBP1 Target | Prioritized Target |
| --- | --- | --- | --- | --- | --- |
| CELECOXIB | NPC | -0.335 | 3.9E-05 | PTGS2 | TRUE |
| MEFENAMIC ACID | HEK293T | -0.276 | 4.0E-03 | PTGS2 | TRUE |
| CHOLIC ACID | NPC | -0.262 | 1.1E-02 | COX4I1 | FALSE |
| CAFFEINE | NEU | -0.245 | 3.4E-02 | PDE4D | TRUE |
| VORINOSTAT | SHSY5Y | -0.244 | 3.6E-02 | HDAC9 | TRUE |
| VALPROIC ACID | NPC | -0.236 | 5.6E-02 | HDAC9 | TRUE |
| CAFFEINE | NPC | -0.224 | 9.8E-02 | PDE4D | TRUE |
| NITRENDIPINE | HEK293 | -0.210 | 1.6E-01 | CACNB2 | FALSE |
| DIFLUNISAL | NPC | -0.198 | 2.2E-01 | PTGS2 | TRUE |
| CRIZOTINIB | NPC | -0.196 | 2.3E-01 | MAP4K1 | TRUE |
| INDOPROFEN | HEK293 | -0.190 | 2.6E-01 | PTGS2 | TRUE |

*PTGS2* (COX-2) emerged as a critical intersection point, targeted by multiple drugs, most notably celecoxib. Previous studies have associated upregulated *PTGS2* with ASD phenotypes and linked its byproduct, PGE2, to the Wnt signaling pathway^85^. Celecoxib ranked as our second-highest-scoring drug for signature reversion (**Figure 3B**) and has previously been shown to reduce dopaminergic dysfunction and improve sensorimotor behavior in neuroinflammatory models^86^, warranting prioritizing for *in vivo* validation in *SETBP1*-HD.

## Discussion

By employing a convergent genomic approach, we identified FDA-approved candidates with the potential to mitigate *SETBP1*-HD phenotypes by targeting their underlying molecular drivers. Mapping our regulatory targets to developmental expression profiles revealed a critical window of target gene expression enrichment early in prenatal development (8-12 pcw), aligning with rapid NPC expansion, followed by a sharp depletion peaking in early childhood (2-5 years), a period of intense synaptic pruning and circuit maturation (**Figure 1**) ^43,44^. Antonyan et al. proposed that SETBP1 may oligomerize, forming nuclear bodies that work with the nuclear lamina, to coordinate nuclear envelope re-assembly and 3D chromatin organization during the cell cycle^6^. In this case, deficiency of SETBP1 would disrupt nuclear breakdown and reassembly, delaying cell cycle exit and pushing NPCs to favor prolonged proliferation over maturation^7,21,49^. This cascade ultimately distorts cortical lamination and circuitry and delays subsequent developmental processes such as myelination^7,44^.

Importantly, the late prenatal shift from SETBP1 target enrichment to depletion marks a critical transition toward synaptic to maturation, differentiation, and synaptic pruning that persists long past birth (**Figure 1**). This suggests that postnatal therapeutic intervention may be a viable strategy to mitigate disease progression^87,88^, similar to recent successful FDA approvals in neurodevelopmental disorders such as Trofinetide for Rett syndrome^89,90^ and Ganaxolone for Cyclin-dependent kinase-like 5 (CDKL5) deficiency disorder (CDD)^88,91^.

Our consensus signature was enriched for pathways driving this pathogenesis, including ‘neuron projection guidance’^6^, ‘regulation of neuron differentiation’^21,50^, ‘neural precursor cell proliferation’^21^, and ‘telencephalon cell migration/development’^21^(**Figure 2A**). Our top signature genes provide mechanistic links to these developmental disruptions. To our knowledge, we are the first to report increased expression of *LIN28* and *SPON1* across *SETBP1*-HD contexts. *LIN28* is a key regulator of developmental timing that suppresses differentiation by preventing let-7 microRNA biogenesis^92,93^. Its upregulation may cause cells to stall in cell cycle, directly contributing to the delayed neurogenesis and extended NPC proliferation observed in *SETBP1*-deficient models^11,21^. Conversely, *SPON1* (F-spondin) plays a critical role in neural development through axon guidance by inhibiting motor axon outgrowth^94^. Its upregulation likely contributes to the peripheral motor phenotypes hypotonia and muscle weakness in *SETBP1*-HD patients^95^. Furthermore, given the recent identification of *SETBP1* as a major driver of Motor Speech Disorders (MSD)^96^, increased *SPON1* expression may mechanistically underpin the fine motor planning deficits distinct to childhood apraxia in *SETBP1*-HD.

Downregulated targets in our consensus signature included *UTY* and *ABAT*. Downregulation of *UTY*, a ubiquitination gene, aligns with recent findings by Wong et al.^11^, who identified decreased ubiquitination as a key *SETBP1* loss-of-function mechanism. Deficiency of another downregulated gene, *ABAT* (a key enzyme in GABA catabolism), is associated with phenotypes overlapping significantly with *SETBP1*-HD, including hypotonia, epilepsy, and developmental delay^4,6,50,97,98^. This downregulation provides a mechanistic link between early transcriptional disruptions and the subsequent destabilization of GABAergic networks.

Querying our consensus signature against the LINCS drug-perturbation database prioritized many neurotransmitter modulators (**Figure 3A**). During fetal development, neurotransmitter systems form the preliminary architectural framework of cortical circuits^44^. GABA dysfunction, in particular, was captured in our consensus signature (*ABAT* downregulation and **Figure 2A**) and has been implicated in ASD^99^.

Importantly, as a potential key therapeutic window in early childhood (**Figure 1**), we also identified therapeutics with established pediatric safety profiles (**Additional file 6**), including buspirone and celecoxib^83,100,101^. Buspirone, an 5-HT1a receptor partial agonist, has successfully been used to treat anxiety and cognitive defects, including in rare disease^83^. Buspirone activates the PI3K-AKT synaptogenesis axis^84^, offering a direct molecular rescue for the hypo-active AKT observed in severely SETBP1-deficient NPCs^6^. As a potent neuromodulator, it chemically stabilizes vulnerable serotonergic circuits and acts as a dopamine D2 autoreceptor antagonist to increase dopamine and noradrenaline levels, which may address overlapping motor and attention deficits^102,103^. Furthermore, buspirone can exhibit anti-inflammatory properties, potentially mitigating the neuroimmune abnormalities often seen in ASD^102^. This provides a strong, multi-modal mechanistic rationale for buspirone potentially restoring synaptic processes, behavioral phenotypes, and cognitive function.

Concurrently, our parallel genomic approaches identified celecoxib, a COX-2 inhibitor. *PTGS2* (COX-2) emerged as a prioritized SETBP1 regulatory target **(Table 2).** Excess COX-2 activity drives the overproduction of PGE2, a known activator of the Wnt/β-catenin pathway^85^. Interestingly, Wnt signaling activity in the context of SETBP1 deficiency models appears to be highly context-dependent. Cardo et al. found downregulated *FOXG1* and upregulation of WNT ligand genes–suggesting hyper-active Wnt/β-catenin activity–in their *SETBP1*-KO NPCs and neurons. Conversely, Antonyan et al. observed decreased β-catenin–suggesting decreased Wnt/β-catenin activity–in *SETBP1*-KO and *SETBP1*-HD-patient-derived iPSC-induced NPCs. We similarly observed conflicting directionality in Wnt-associated DEGs (e.g., *FZD6* and *ZNRF3*) when comparing Cardo and Shaw data sets^21,50^ (**Additional file 9**). These inconsistencies highlight the need for further research to characterize the role of Wnt signalling across *SETBP1*-HD contexts. Because *PTGS2* upregulation is also strongly associated with neuroinflammation and excitotoxicity in ASD^104,105^, celecoxib may offer a dual therapeutic benefit: resolving potential Wnt-driven proliferation defects and mitigating neuroinflammation-associated dendritic pathology.

## Limitations

This study relies primarily on *in silico* consensus modeling derived from stem cells (hESC/iPSCs) and animal transcriptomics. While these platforms provide critical molecular insights, they do not fully capture the systemic complexity of the living human brain. Furthermore, previous studies demonstrate that different *SETBP1* variants produce highly heterogeneous functional and clinical profiles^11,50,106^. Because our consensus approach maximizes consistency to identify broadly applicable mechanisms, it may overlook distinct patient subgroups or variant-specific therapeutic targets. Finally, although our developmental analysis highlights a critical window for early intervention, prenatal administration of COX-2 inhibitors like celecoxib carries significant fetal risks (**Additional file 6**). Therefore, drug candidates identified here require rigorous *in vivo* validation and clinical trials to establish safety and efficacy.

## Conclusion

This study establishes a comprehensive SETBP1 regulatory target list and a robust, consensus transcriptomic signature for *SETBP1* haploinsufficiency, resolving inconsistencies across heterogeneous disease models. We identify *LIN28A* and *SPON1* as novel, convergent drivers of the *SETBP1*-HD pathology: *LIN28A* upregulation mechanistically underpins the delayed neurogenesis and proliferative stalling, while *SPON1* overexpression provides a molecular basis for the motor and speech apraxia phenotypes via inhibition of motor axon guidance. By integrating these molecular insights with inversely-associated drug perturbation signatures, we prioritized celecoxib and buspirone as repurposable candidates. These drugs offer a dual therapeutic strategy, respectively. Ultimately, this genomics-driven approach provides actionable, context-aware therapeutic avenues for immediate investigation in *in vivo* models and clinical settings.

## Additional Files

### Additional file 1: Graphical abstract

Additionalfile1.png

Graphical abstract of *SETBP1-*HD drug repurposing schema used in this study. Created in BioRender. Wilk, Elizabeth. (2026) https://BioRender.com/5m6vkkm

### Additional file 2: Full SETBP1 targets table

Additionalfile2.xlsx

Table of comprehension SETBP1 targets with gene/protein name (target), synonymous target names, organism, interaction type with SETBP1, regulatory effect by SETBP1, evidence (score parameters, SETBP1 mass spectrometry IP from Antonyan et al., and notes), sources (databases and publications), and Pharos data (Target Development Level, Family, and Novelty).

### Additional file 3: Data sets used in SETBP1-HD analyses

Additionalfile3.xlsx

Table describing samples used in these analyses including study name used in manuscript, dataset accession, number of samples used, and model details (organism, cell types, and genotypes).

### Additional file 4: PCA of data sets combined

Additionalfile4.pdf

EDA of data sets: PC1 splits Tanaka (only mouse) vs rest of data sets, PC2 splits mostly by cell type, clustering by data set A) PC1 and PC2, colored by cell type (purple for human, yellow for mouse) and shape indicating data set by first author name (circle = Cardo, triangle = Shaw, square = Tanaka). B) PC2 and PC3, colored by cell type (purple = HSC, blue = iPSC, green = Neuron, yellow = NPC) and shape indicating genotype (circle for control, triangle for SETBP1-HD).

### Additional file 5: Spatio-temporal SETBP1 target expression sex differences

Additionalfile5.xlsx

Table of raw and FDR-adjusted p-values from Wilcoxon rank sum test comparing SETBP1 target expression by sex at each GTEx age range for each tissue.

### Additional file 6: Full, Annotated Drug Candidates table

Additionalfile6.xlsx

Drug candidates filtered for 0 < WTCS signatures and neurologically-representative cells from signatureSearch outputs, signatureSearch annotations, SETBP1 targets as drug targets, and DailyMed extracted drug label data for black box warnings (BBW), pediatric indication, pregnancy and lactation risk summary, pharmacological kinetics, and blood brain barrier (BBB) information.

### Additional file 7: DEGs by Contrast and Data Set

Additionalfile7.png

Barplots indicating number of DEGs by individual contrasts for Cardo (purple) and Shaw (green) data sets.

### Additional file 8: Count of DEGs for signature methods, colored by geneset source

Additionalfile8.png

Upset plot comparing DEGs across contrasts in Shaw (blue) and Cardo (yellow) data, ordered by overlap comparison (i.e., from DEGs unique to contrasts to DEGs shared across every contrast). “Set size” indicates the full set of DEGs by contrast, “Intersection size” indicates the number of DEGs in a given overlap, and points/lines indicate the particular overlap comparison.

### Additional file 9: LogFCs of Genes from RRA Excluded for Low Effect Size and/or Inconsistent Direction

Additionalfile9.pdf

Heatmap displaying RRA DEGs that were filtered from consensus signature for having meta-analysis logFCs (|logFC| <= 0.1) due to low effect size and/or inconsistent effect directionality.

## Declarations

### Ethics approval and consent to participate

Consent for publication

### Availability of Data and Materials

The data used for this study are publicly available on GEO and SRA (Cardo: PRJNA746989, Shaw: PRJNA1093043, and Tanaka PRJNA898600 and PRJNA974068), from Wong et al. supplementary information (DEGs from SourceData6_forFIg5c_DEG.xlsx and SourceData10_forFig7b_DEG, metadata from Supplementary Data 5; 41467_2025_64074_MOESM7_ESM.xlsx and Supplementary Data 7; 41467_2025_64074_MOESM9_ESM.xlsx for patient-derived fibroblasts and induced neurons, respectively), from the GTEx Portal (https://gtexportal.org/home/downloads/adult-gtex/overview, GTEx 2025-08-22 v11 RNASeQCv2.4.3 TPM and metadata files [v11 Sample and Subject Attributes DS] and dGTEx 2025-07-21 v1 RNASeQCv2.4.3 TPM and metadata files [v1 2026-01-30 Sample and Subject Attributes]), BrainSpan online atlas (https://www.brainspan.org/static/download.html, api v2, developmental transcriptome zip file with expression matrix, row metadata [genes], column metadata [samples]), All code to replicate analyses in this study are available on GitHub https://github.com/lasseignelab/setbp1_hd and Zenodo (10.5281/zenodo.20073553)^79^ and was built using the CAPTURE framework (cap version 1.0)^77^. Docker images used to execute analyses are available on DockerHub (ncbi/sra-tools:3.2.1, projectassistant/sitemap-scraper:latest, lizzyr/lw_models:0.1.1, and lizzyr/lw_sigsearch:1.0.0) and Zenodo (10.5281/zenodo.20071673)^78^. Intermediary and final outputs are available on Zenodo (10.5281/zenodo.20073644)^73^).

### Competing Interests

The authors have no competing interests to declare.

### Funding

We are grateful to our funders for this work: the SETBP1 Society (2024 Microgrant) and the UAB Pilot Center for Precision Animal Modeling (C-PAM) (U54-OD030167)

### Authors’ Contributions

Project was conceptualized by BNL and EJW; data curation by EJW and ST; formal analysis by EJW and ST; funding acquisition by BNL; investigation by EJW and ST; methodology by EJW; project was supervised by BNL and EJW; original code for analysis was written by EJW and independently validated by TMS and ST; original manuscript drafted by EJW and ST; draft editing by EJW, ST, TMS, and BNL.

## Acknowledgements

We thank the Lasseigne Lab members for their thoughtful input and discussion. We also extend our deep gratitude to the SETBP1 Society and the broader community of researchers, clinicians, patients, and caregivers. Your dedication, open sharing of data, and collaborative discussions have been instrumental in advancing this research and improving the care of those affected by *SETBP1* variation.

## Notes

https://zenodo.org/records/20073644

https://zenodo.org/records/20073553

https://zenodo.org/records/20071673

### Competing Interest Statement

The authors have declared no competing interest.

