## Supplementary figures and images for "Intersection of Regulatory Analysis and Signature Reversion Uncovers Therapeutic Drugs and Targets for *SETBP1*-HD"

### Additional file 1

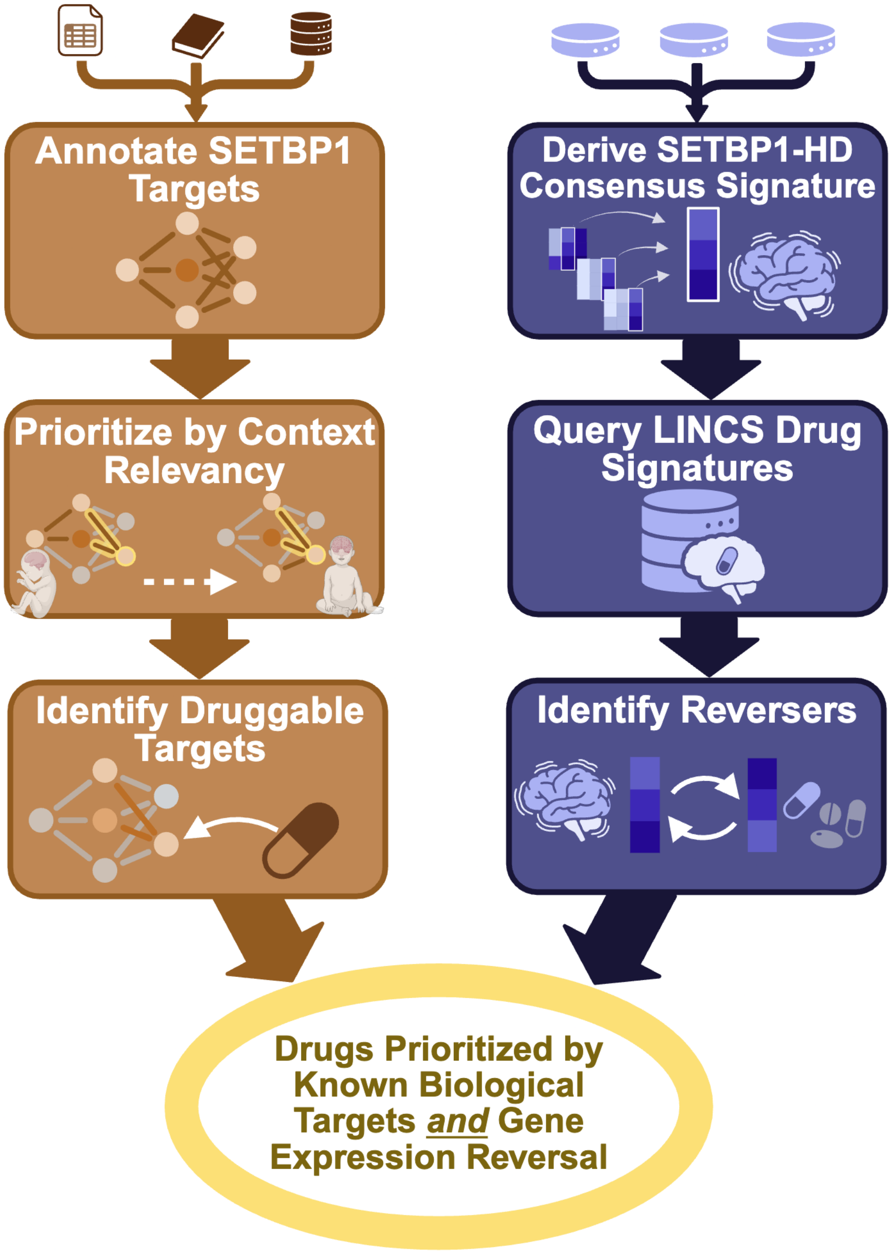

### Additional file 4

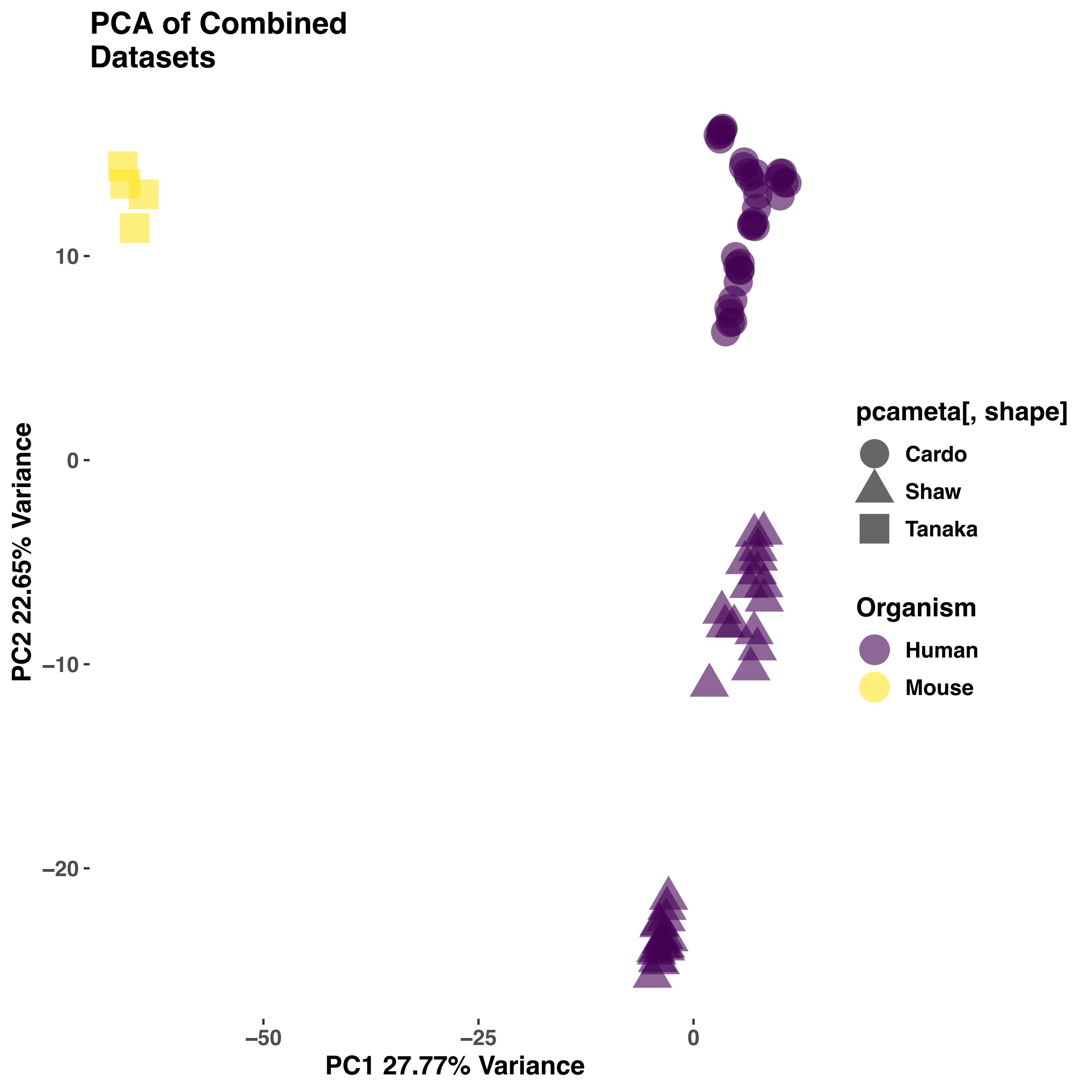

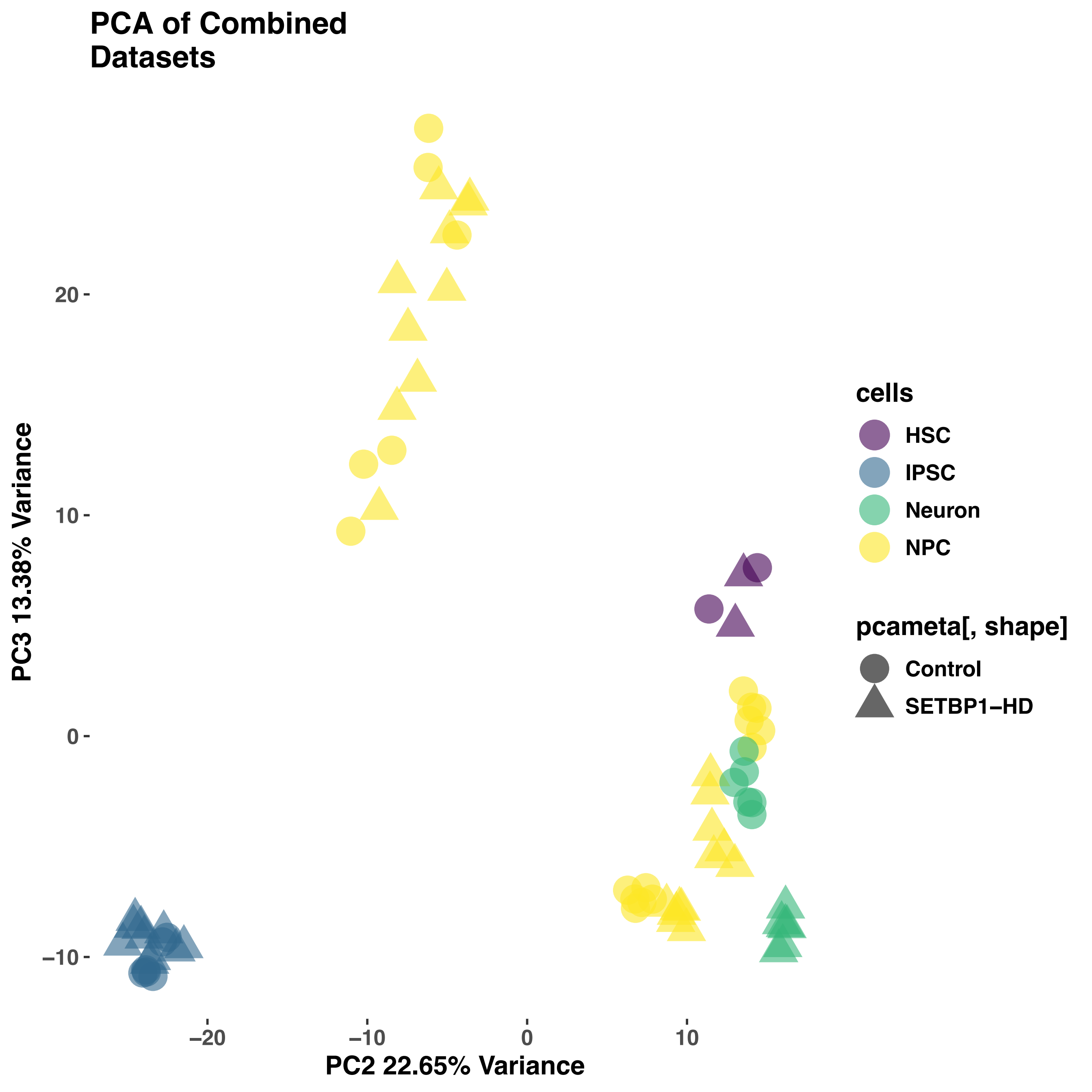

### Additional file 7

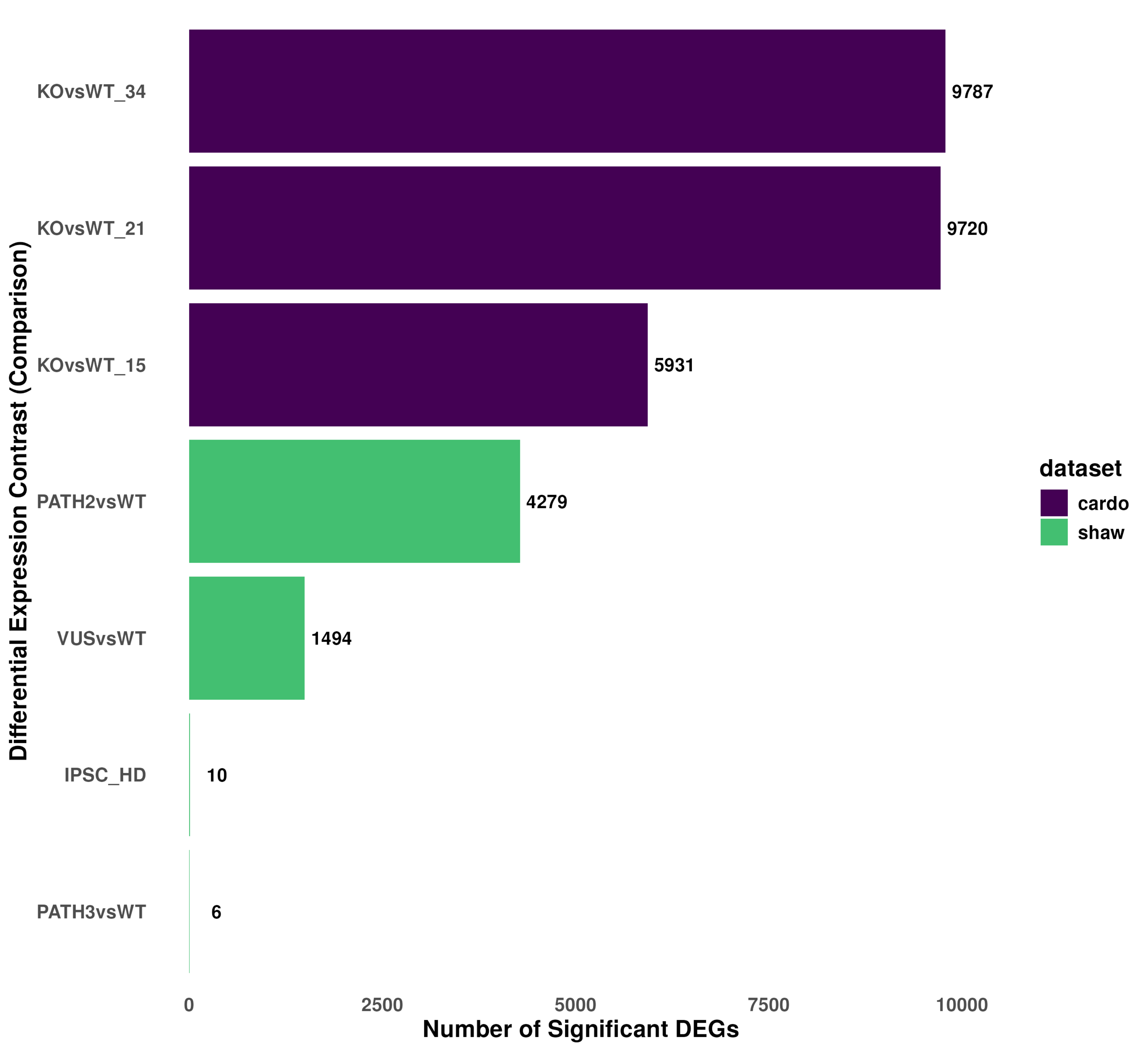

### Additional file 8

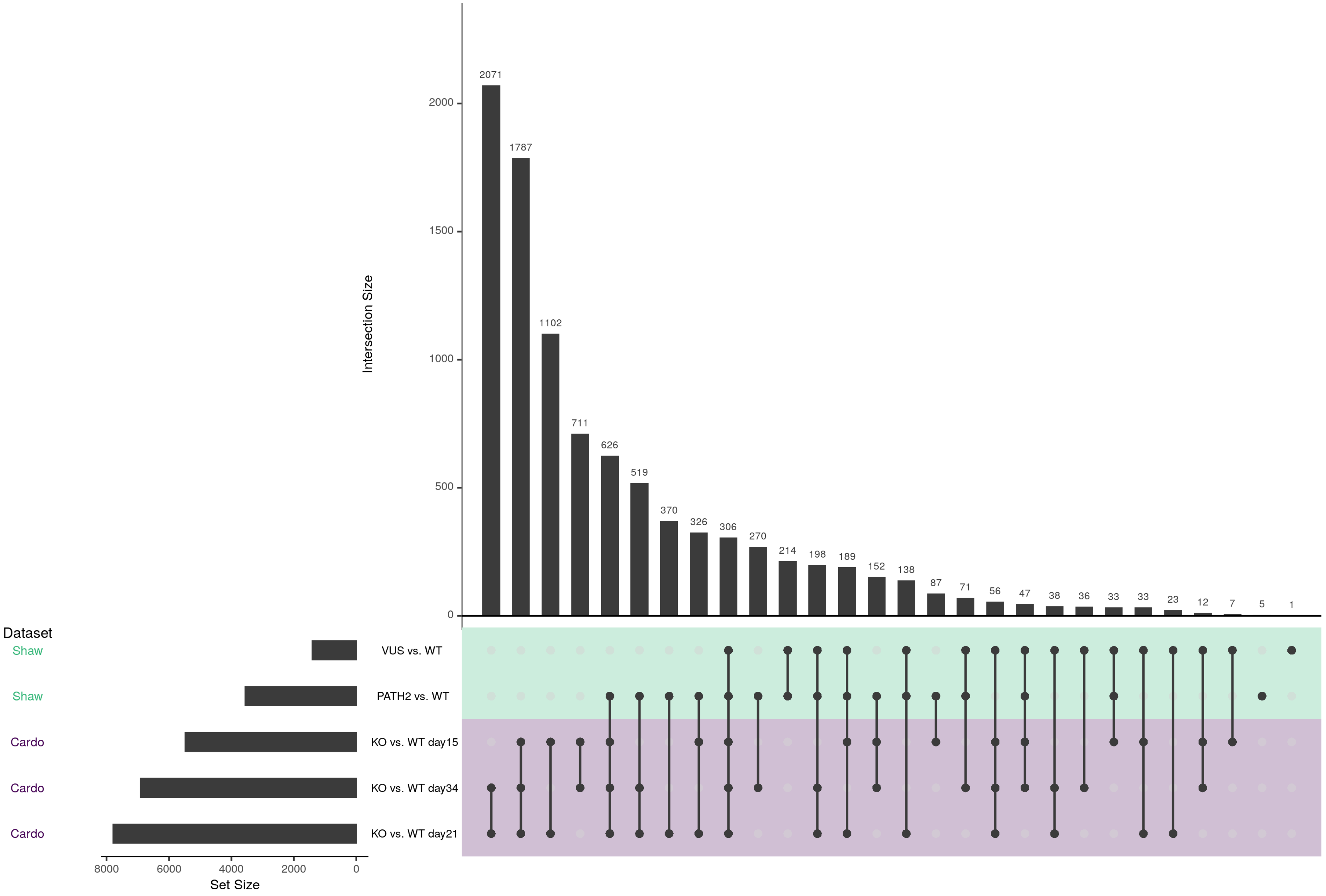

### Additional file 9

RRA Genes Excluded by Meta logFC Threshold−Based SETBP1−HD Consensus Signature

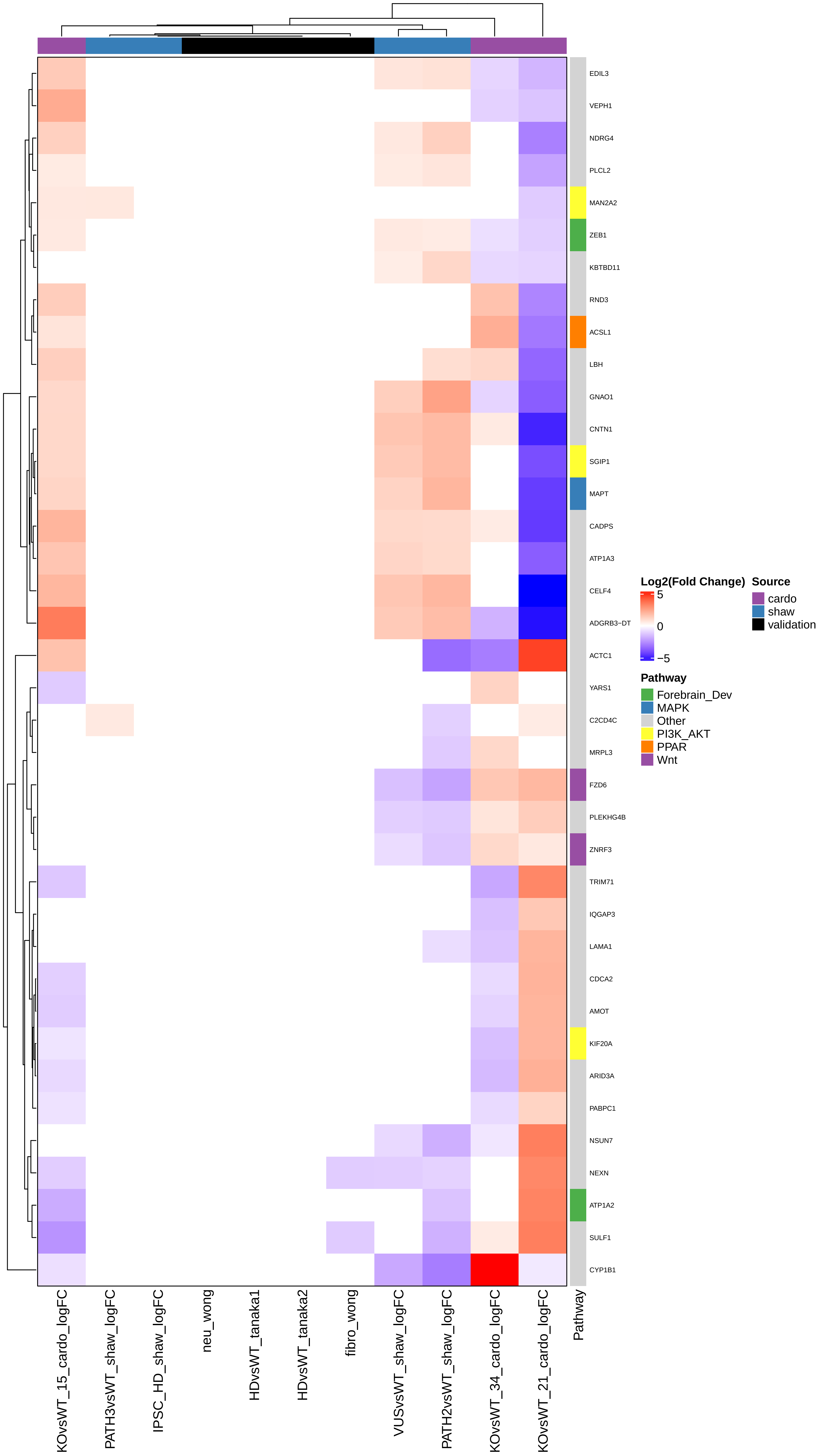
